# Analysis of mouse lens morphological and proteomic abnormalities following depletion of βB3-crystallin

**DOI:** 10.1101/2024.12.30.630781

**Authors:** Danielle Rayêe, Phillip A. Wilmarth, Judy K. VanSlyke, Keith Zientek, Ashok P. Reddy, Linda S. Musil, Larry L. David, Ales Cvekl

**Affiliations:** Departments of Ophthalmology and Visual Sciences and Genetics, Albert Einstein College of Medicine, Bronx, New York 10461; Proteomics Shared Resource, Oregon Health Sciences University, 3181 Southwest Sam Jackson Park Road, Portland, OR 97239; Department of Chemical Physiology and Biochemistry, Oregon Health & Science University, 3181 Southwest Sam Jackson Park Road, Portland, OR, 97239

**Keywords:** βB3-crystallin, cataract, cytoplasm, lens, proteomes, isobaric labeling, quantitative proteomics

## Abstract

Crystallin proteins serve as both essential structural and as well as protective components of the ocular lens and are required for the transparency and light refraction properties of the organ. The mouse lens crystallin proteome is represented by αA-, αB-, βA1-, βA2-, βA3-, βA4-, βB1-, βB2-, βB3-, γA-, γB-, γC-, γD-, γE, γF-, γN-, and γS-crystallin proteins encoded by 16 genes. Their mutations are responsible for lens opacification and early onset cataract formation. While many cataract-causing missense and nonsense mutations are known for these proteins, including the human CRYBB3 gene, the mammalian loss-of function model of the *Crybb3* gene remains to be established. Herein, we generated the first mouse model via deletion of the Crybb3 promoter that abolished expression of the βB3-crystallin. Histological analysis of lens morphology using newborn βB3-crystallin-deficient lenses revealed disrupted lens morphology with early-onset phenotypic variability. In-depth lens proteomics at four time points (newborn, 3-weeks, 6-weeks, and 3-months) showed both down- and up-regulation of various proteins, with the highest divergence from control mice observed in 3-months lenses. Apart from the βB3-crystallin, another protein Smarcc1/Baf155 was down-regulated in all four samples. In addition, downregulation of Hspe1, Pdlim1, Ast/Got, Lsm7, Ddx23, and Acad11 was found in three time points. Finally, we show that the βB3-crystallin promoter region, which contains multiple binding sites for the transcription factors AP-2α, c-Jun, c-Maf, Etv5, and Pax6 is activated by FGF2 in primary lens cell culture experiments. Together, these studies establish the mouse *Crybb3* loss-of-function model and its disrupted crystallin and non-crystallin proteomes.

## INTRODUCTION

The ocular lens is a cellular tissue comprised from two compartments, the anterior lens epithelium and posterior lens fiber cells, which represents bulk of the tissue (see Bassnett et al., 2011; McAvoy et al., 2017). Lens transparency and light refraction require high concentrations of lens-preferred crystallin proteins that accumulate to concentrations as high as 450 mg/ml in the central region of the fiber cell compartment (“lens nucleus”) (see Bloemendal et al., 2004; Bassnett et al., 2011). The maintenance of lens proteostasis over the organismal lifespan is critical for normal lens function and its disruption results in cataract formation. Lens crystallins are classified as α- and β-/γ-families of proteins and are related to small heat shock proteins (Franck et al., 2004) and Ca^2+^-binding microbial proteins, respectively (Kappe et al. 2010; Srivastava et al. 2014). Mutations in crystallin genes cause congenital and/or early childhood cataracts (see Shiels and Hejtmancik, 2021) while aging-driven changes in crystallin proteins such as racemizations, deamidation, amino acid posttranslational modifications, and protein truncations are found in age-related cataract lenses (see Bloemendal, et al., 2004; Lampi et al., 2014; Quinlan and Clark, 2022; Cvekl and Vijg, 2024).

The βA1-, βA2-, βA3-, βA4-, βB1-, βB2-, βB3-, γA-, γB-, γC-, γD-, γE, γF-, γN-, and γS-individual crystallin genes arose from ancestral gene duplication events and form two subfamilies of β- and γ-crystallins (Lampi et al., 2014; Cvekl et al., 2017). Their structures involve a pair of N- and C-terminal Greek key domains that are highly hydrophobic and rich in aromatic and sulfur-containing amino acid residues (Serebryany and King, 2014). The mouse and human βB3-crystallin are 211 amino acid long proteins (MW 24.2 kDa) encoded by the *Crybb3* (mouse) and *CRYBB3* (human) genes. These genes are comprised of five exons, located on chromosome 5 (mouse) and 22 (human). Human and mouse βB3-crystallin represent approximately 1% and 8% of newborn human and mouse lens water-soluble proteins, respectively, and decrease in abundance during aging (Ueda et al., 2002; Halverson-Kolkind et al., 2024). Several cataract-causing heterozygous mutations in the βB3-crystallin proteins have been reported, including the G165R (Riazuddin et al., 2005; Jiao et al., 2016), V194G (Reis et al., 2013), G76R, R105Q, and G165R missense mutations (Li et al., 2016). Microduplication of 495 kb subregion originating from the 22q11.23 including adjacent CRYBB2 and CRYBB3 genes also caused bilateral cataracts (Trkova et al., 2016). Analysis of bovine lenses identified a 15 amino acid long peptide (residues 153 to 167) originating from the βB3-crystallin generated after oxidation and proteolysis (Senthilkumar et al., 2002), and follow-up biochemical studies showed that this fragment binds to βB2-crystallins and enhances their aggregation, precipitation, and light scattering (Udupa and Sharma, 2005a; Udupa and Sharma, 2005b). Analysis of another member of this subfamily, i.e., the βB1-crystallin, focused on the role of N-terminal truncation, liquid-liquid phase separation, and mechanisms of cataractogenesis (Annunziata et al., 2005). It has been shown recently that C-terminally truncated βB1-crystallin (Y204X) cataract-causing mutation allows dimer formation with different surfaces leading to their aggregation and protein degradation (Jing et al., 2023).

Pioneering crystallin gene loss-of-function studies initially focused on the mouse *Cryaa* and *Cryab* models encoding the αA- and αB-crystallins, respectively. Somatic depletion of the αA-crystallin proteins resulted in smaller lenses and their opacifications were initiated within the lens “nucleus”. This depletion caused the αB-crystallin proteins to be recovered in the insoluble cytoplasmic fraction (Brady et al., 1997). In contrast, the αB-crystallin depleted lenses developed normally with all other crystallins found close to their normal levels (Brady et al., 2001). Finally, the double somatic knockout model (*Cryaa*^-/-^; *Cryab*^-/-^) produced significantly smaller lenses with many morphological abnormalities (Boyle et al., 2003). A spontaneous *Nuc1* mutation exists in the rat *Cryba1* gene encoding both βA1- and βA3-crystallins (see Zigler and Sinha, 2015). The heterozygous *Nuc1* (*Cryba1*) mutation causes nuclear cataracts while the homozygous mutation leads to microphthalmia and early disruption of lens structure as well as retinal abnormalities (Sinha et al., 2005). An unexpected retention of nuclei was observed in the presumptive organelle-free zone of the lens (Sinha et al., 2005) indicating that βA1- and βA3-crystallins also play a role in lens fiber differentiation and denucleation (see Zigler and Sinha, 2015). Finally, depletion of mouse βB2-crystallins generates normal sized lenses with cortical cataracts developing after several months (Zhang et al., 2008). Interestingly, the βB2-crystallin also plays major roles in the retina, brain, ovaries, and testes (Gao et al., 2014; Heermann et al., 2019; Li et al., 2021; Bauer et al., 2024).

Based on these earlier studies, we hypothesized that somatic depletion of the βB3-crystallin proteins would perturb lens proteostasis, possibly affecting early lens morphogenesis, while not resulting in any major abnormalities outside of the eye. To generate new insights into the function of βB3-crystallin, we generated a loss-of-function model by deleting its lens-specific promoter region (Zhao et al., 2019a) using the CRISPR-Cas9 technology. Earlier studies of βA1-, βB1-, and βB2-crystallin genes (*Cryba1*, *Crybb1*, and *Crybb2*) identified lens-specific promoters within a range of ∼170 to 460 bp (Chambers et al., 1995; Duncan et al., 1996; McDermott et al., 1996). Our previous studies identified Pax6-binding sites and “open” chromatin in the mouse *Crybb3* promoter (Sun et al., 2015; Zhao et al. 2019a). Thus, deletion of the promoter region was a simple option to generate the *Crybb3* null allele model. Depletion of expression of the βB3-crystallin protein resulted in variable lens abnormalities frequently detectable in homozygous newborn eyes while heterozygous eyes appeared consistently normal. Consequently, a major portion of this study involved characterization of control and mutated lens proteomes and their comparison at different developmental ages. Our results also show that βB3-crystallin loss-of-function alters the expression of BAF-complex subunit/Baf155, 10 kDa heat shock protein (Hspe1), PDZ and LIM domain protein 1/elfin (Pdlim1), aspartate aminotransferase (Ast/Got), U6 snRNA-associated Sm-like protein LSm7 (Lsm7), RNA helicase (Ddx23), and Acyl-CoA dehydrogenase family member (Acad11) in at least three of four-time comparisons.

## MATERIALS AND METHODS

### βB3-crystallin promoter excision mouse line production

All experiments and animal husbandry were conducted following the approved protocol of the Institutional Animal care and Use Committee at Albert Einstein College of Medicine and the ARVO Statement for the Use of Animals in Ophthalmology Research. The mouse line mutants were generated using CRISPR/Cas9-mediated genome editing. A pair of guide RNAs targeting the upstream (targeting sequence: GAGGCCACATCAGTTACTTGGGG) and the downstream region (targeting sequence: CCCTTGTCTGGCTCACTCGAGGG) of the βB3-crystallin promoter were designed by an online software (www.benchline.com) and generated by *in vitro* transcription. Mice were genotyped by PCR using tail DNAs and a pair of primers to identify the expected target sequence (Foward: 5’-GCCTCTTCTTCTGGCATCT-3’; Reverse: 5’-CCAGCAGCTTCTGCTCTGTA-3’).

### Hematoxylin and Eosin Staining (H&E) and Immunohistology

Mouse heads (newborns) were collected and fixed in PFA 4% overnight at 4°C and cryoprotected in 30% sucrose solution (s/H_2_0) for 24 hours. The tissue was embedded in optimal cutting temperature (OCT) and kept at -80 °C until sectioning. Serial sections of 10 µm thickness through the optic nerve were taken and stained with HE following manufacturer instructions (Hematoxylin and Eosin Stain kit, Abcam ab245880) for morphological analysis. We performed immunofluorescence staining as described elsewhere (Limi et al., 2018). Briefly, tissue sections were incubated for 10 minutes at 90°C in Reveal Decloaker solution (RV1000MRU, Biocare Medical), blocked in 1% BSA for 1 hour at room temperature and incubated overnight at 4°C with primary antibodies (βB3-crystallin Antibody; Santa Cruz sc-374374) diluted in blocking solution. Slides were washed three times in PBS and incubated with anti-mouse secondary antibody for 2h at room temperature (Alexa Fluor 594, Jackson Labs 115-585-003) and mounted with mounting media (Vectashield).

### Western blotting of βB3-crystallin

Western blot analysis of βB3-crystallin was conducted as previously described. Blots were incubated overnight at 4°C with primary antibodies against βB3-crystallin (Santa Cruz; sc-374374) and β-actin (Santa Cruz; sc-sc-47778), after which HRP-conjugated antibodies (Cell Signaling Anti-Rabbit IgG; 7074P2 and Cell Signaling Anti-Mouse IgG; 7076S) were used for detection by enhanced chemiluminometry detection (ECL). The membrane was scanned following manufacturer instructions (ChemiDoc imaging system, GelDoc2000 Software; Bio-Rad).

### Quantitative Real-Time PCR

Trizol was used to extract RNA from 10 whole lenses of WT, heterozygous and Crybb3 promoter deleted mice to quantify Crybb3 (5’-CAGCCGACGTAGTGACATTC-3’; 5’-ACTGAAGGCTGGGTTCTCAA-3’) mRNA levels. 5 µg of the RNA extracted was DNAse treated (Quiagen) and cDNA was synthesized using the SuperScript III First-Strand Synthesis System (Thermo Fisher). SYBR Green Master Mix (Invitrogen) was used to perform qPCR on the Applied Biosystems QuantStudio3 system. GAPDH (5’-ATTGCCCTCAACGACCACT-3’; 5’-ATGAGGTCCACCACCCTGT-3’) was used as a housekeeping gene control to normalize relative gene expression. Experiments were made in triplicates and statistical analysis were performed using t -Test with Bonferroni correction.

### Proteomics samples

Whole lenses from wild type and knock out newborn, 3-weeks, 6-weeks and 3-month-old mice were dissected and fresh-frozen as previously described (Zhao et al., 2019b). Lenses were pooled from 20, 15, 5, and 5 mice at the respective ages and two biological replicates for WT and KO at each age were prepared (16 samples in total). Lenses were stored at -80^0^C and thawed at 2-5^0^C before processing. To each tube of lenses, 300 µL of 67 mM TEAB was added, centrifuged at 100xg for 3 min, and samples were transferred to clean 1.5 mL LoBind Tubes. 100 uL of 20% SDS was added to each tube along with 40-50 mg of acid washed glass beads (Sigma PN#G8772-10G). Final buffer composition was 5% SDS, 50 mM TEAB in 400 µL. A Bertin Precellys Evolution Bead Beater with Cryolys attachment was used for the initial homogenization step: 6 x 20 sec cycles (with a 30 sec pause between cycles), 6300 rpm, at 4C. Samples were then heated with shaking at 70^0^C for 10 min and centrifuged for 3 min at 8000xg. Finally, samples were sonicated in a Diagenode Bioruptor Pico (30 sec on, 30 sec off for 10 cycles). Protein content was determined using a Pierce BCA protein assay kit (Thermo Scientific Catlg #23227).

### Protein tryptic digestion

A volume equal to 50 µg of protein from each sample was transferred to 1.5 ml Lobind centrifuge tubes, and SDS protein extraction buffer (5% SDS, 50mM TEAB, pH8) was added to bring the final volume to 75 µl. Samples were reduced by adding 3.4 µl of 0.5M dithiothreitol and incubated at 95°C for 10 min. Samples were alkylated by the addition of 6.8 µl of 0.5M iodoacetamide and incubation at room temperature for 30 min in the dark. Samples were then acidified by the addition of 8.5 µl of 12% phosphoric acid, and 562 µl of SDS protein extraction buffer (90% aqueous methanol, 100 mM TEAB, pH 8) was added. Samples were transferred 165 µl at a time to S-trap micro columns (Protifi, Farmingdale, NY) inserted into 1.5 ml of polypropylene tubes. Samples were centrifuged at 4,000 x g for 3 min between each addition of the sample. S-trap columns were washed 6x using 160 µl 90% methanol, 100 mM TEAB followed by centrifugation at 4,000 x g for 3 min between each wash step. S-trap columns containing the bound sample proteins were transferred to 1.5 ml Lobind centrifuge tubes and 40 µl of 80 ng/µl sequencing grade modified trypsin (Promega, Fitchburg, WI, Cat # V5111) in 50 mM TEAB was added. S-trap columns were capped, and digestion was performed at 37°C overnight in a humidified chamber.

After digestion, peptides were eluted by sequential addition of 40 µl of 50 mM TEAB, 40 µl of 0.2% aqueous formic acid, and 40 µl of 50% acetonitrile, 0.2% formic acid, with centrifugation at 4,000 x g for 4 min between each addition of elution buffer. The eluted fractions were pooled and then dried by vacuum centrifugation, 100 µl of 50% methanol added, and samples again dried. Samples were reconstituted with 100 µl of water at 37°C in a shaker and peptide concentrations determined using a Pierce Peptide Assay (ThermoFisher Scientific, Cat # 23225). For the TMTpro labeling step described below, 15 µg of peptide digest from each sample was dried by vacuum centrifugation. Dried peptides were reconstituted by adding 20 µL of 100 mM TEAB and shaking at 37°C for 15 minutes.

### TMTpro labeling and normalization run

A tandem mass tag (TMTpro) 16-plex reagent kit (ThermoFisher Scientific, Cat # 90309) was used to label the digested peptide samples. Each TMTpro 16-plex reagent (200 µg) dissolved in 12 µl of anhydrous acetonitrile was added to 15 µg of peptide sample in 20 µl of 100mM TEAB, and labeling performed by shaking at room temp for 1h. After the incubation, 2 µl of each labeled peptide digest were combined; 2 µl of 5% hydroxylamine added, and samples incubated at room temp for 15 min, then dried by vacuum centrifugation. The remaining volume of each labeled sample was frozen at -80°C without hydroxylamine addition (in case relabeling was required).

The 2 µl of each combined TMTpro labeled sample was then dissolved in 20 µl of 5% formic acid and 2 µg of peptides analyzed by a single 140 min LC-MS/MS method using an Orbitrap Fusion Mass Spectrometer to check labeling efficiency (typically >90%) and determine the volume of each sample to provide similar total reporter ion intensities in the final combined sample for the 2D-LC/MS analysis. The TMTpro labeled samples were thawed and aliquots were removed, based on the calculated normalization factors that would produce 50 µg of total peptides from all 16 samples. To quench the labeling reaction, 5% hydroxylamine was added to bring the total hydroxylamine concentration to 0.5% followed by incubation for 15 min at room temperature. The remaining labeled peptides not used for the 2D-LC/MS were stored at -80°C in case a second run was needed.

### Two-dimensional liquid chromatography/mass spectrometry

The multiplexed mouse lens samples were dissolved in 10 mM ammonium formate, pH9 buffer and injected onto a NanoEase 5 µm XBridge BEH130 C18 300 µm x 50 mm column (Waters Corporation, Milford, MA) at 3 µl/min in a mobile phase containing 10 mM ammonium formate (pH 9). Peptides were eluted by sequential injection of 20 µl volumes of 17, 20, 21, 22, 23, 24, 25, 26, 27, 28, 29, 30, 31, 32, 33, 34, 35, 40, 50, 90% ACN (20 fractions).

Eluted peptides were diluted at a 3-way union with mobile phase containing 0.1% formic acid at a 24 µl/min flow rate and delivered to an Acclaim PepMap 100 µm x 2 cm NanoViper C18, 5 µm trap (Thermo Fisher Scientific) on a switching valve. After 10 min of loading, the trap column was switched on-line to a PepMap RSLC C18, 2 µm, 75 µm x 25 cm EasySpray column (ThermoFisher Scientific). TMTpro 16-plex labeled peptides were then separated at low pH in the second dimension using a 5–25% ACN gradient over 100 min in the mobile phase containing 0.1% formic acid at a 300 nL/min flow rate. Tandem mass spectrometry data was collected using an Orbitrap Fusion Tribrid instrument configured with an EasySpray NanoSource (Thermo Scientific, San Jose, CA). Survey scans were performed in the Orbitrap mass analyzer at resolution = 120,000, with internal mass calibration enabled, and data-dependent MS2 scans using dynamic exclusion performed in the linear ion trap using collision-induced dissociation. Reporter ion detection was performed in the Orbitrap mass analyzer (resolution = 50,000) using MS3 scans following synchronous precursor isolation of the top 10 fragment ions in the linear ion trap, and higher-energy collisional dissociation in the ion-routing multipole. There were 363,428 MS2/MS3 instrument scans acquired from the 20-fractions.

### Proteomics Data Analysis

The binary instrument files were processed with the PAW pipeline (Wilmarth et al., 2009). Binary files were converted to text files using MSConvert (Chambers et al., 2012). Python scripts extracted TMT reporter ion peak heights and fragment ion spectra in MS2 format (McDonald et al., 2004). The Comet search engine (version 2016.03) (Eng et al., 2013) was used: 1.25 Da monoisotopic peptide mass tolerance, 1.0005 Da monoisotopic fragment ion tolerance, semi tryptic cleavage with up to two missed cleavages, variable oxidation of methionine residues (+15.9949 Da), static alkylation of cysteines (+57.0215 Da), and static modifications for TMTpro labels (+304.2071 Da at peptide N-termini and at lysine residues). Searches used UniProt proteome UP000000589 (Mus musculus, taxon ID 10,090) canonical FASTA sequences (21,957 proteins, downloaded July 2023). Common contaminants (175 sequences excluding any albumins) were added, and sequence-reversed entries were concatenated for a final protein FASTA file of 44,264 sequences.

Top-scoring peptide spectrum matches (PSMs) were filtered to a 1% false discovery rate (FDR) using interactive delta-mass and conditional Peptide-prophet-like linear discriminant function scores (Keller et al., 2002). Incorrect delta-mass and score histogram distributions were estimated using the target/decoy method (Elias and Gygi, 2007). The filtered PSMs were assembled into protein lists using basic and extended parsimony principles and required two distinct peptides per protein. The final list of identified proteins, protein groups, and protein families were used to define unique and shared peptides for quantitative use. Total (summed) reported ion intensities were computed from the PSMs associated with all unique peptides for each protein.

The protein intensity values for each biological sample in each biological condition were compared for differential protein expression using the Bioconductor package edgeR (Robinson et al., 2010). Result tables contained typical proteomics summaries, reporter ion intensities, and statistical results. Jupyter notebook files showing data processing and statistical analysis details for this experiment are available in the ProteomeXchange PRIDE partner archive for this project (dataset identifier: PXD059220) (Perez-Riverol et al., 2022).

### Cell culture and FGF2 assay

The mouse βB3-crystallin promoter fragment (-260/+250) was inserted into the eGFP-1 reporter (GeneScript, Piscataway, NJ) as described earlier in our similar experiments (Xie et al., 2016; Martynova et al., 2018). Two mutants, M1 and M2, include the disruption of one (M1; upstream binding site) or two (M2) Pax6 binding sites (Fig. 1B). Primary embryonic lens epithelial cell cultures were prepared from E10 chick lenses and plated on laminin-coated 96-well plates at a density of 0.85 x 105cells/ well. Cells were cultured without serum in M199 medium plus BOTS (2.5 mg/ml bovine serum albumin, 25 µg/ml ovotransferrin, 300 nM selenium), supplemented with penicillin G and streptomycin. One day after plating, the cultures were transfected with Lipofectamine 2000 (Invitrogen, Waltham, MA, USA) in M199 medium with 0.125 µg (1X) or 0.25µg (2X) reporter plasmid DNA per well. After five hours, cells were cultured in the presence or absence of 10 ng/ml FGF2 (R&D Systems, Minneapolis, MN) for 6 days. EGFP expression in lens cultures was monitored by immunofluorescent microscopy before solubilizing them in SDS-PAGE sample buffer. Equal amounts of lysate were transferred to PVDF membranes that were probed with primary antibodies JL-8 anti-GFP (Clontech, USA) and β-actin, clone C4 (MilliporeSigma, Billerica, MA, USA) and secondary antibody conjugated to Alexa Fluor 680 (Molecular Probes, USA). The staining was analyzed by the Odyssey infrared imaging system (LI-COR Biosciences, USA). The results of EGFP immunoreactivity were normalized to β-actin staining in the same sample as we described earlier (Xie et al., 2016; Martynova et al., 2018).

**Figure 1:**
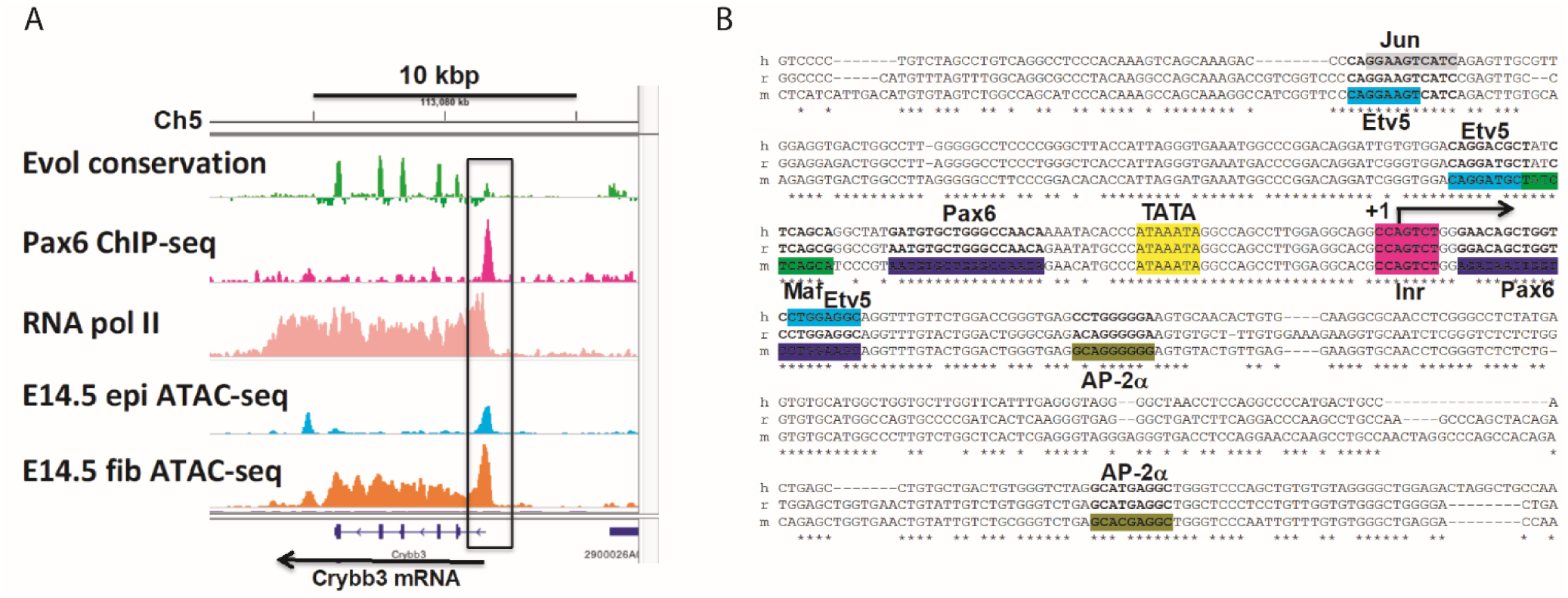
Genomic organization of the mouse Crybb3 gene and its promoter. **A.** Mouse βB3-crystallin gene and its “open” chromatin domains detected by ATAC-seq (Zhao et al. 2019a), Pax6-and RNA Polymerase II binding detected by ChIP-seq (Sun et al. 2015a; Sun et al. 2015b). An 18 kb region of the *Crybb3* locus located on mouse chromosome 5 (ch5) and direction of transcription (arrow in the bottom) are shown. Evolutionary conservation tracks, Pax6 binding peak, RNA Polymerase II, open chromatin in epithelial and fiber cells detected by ATAC-seq are shown in green, purple, light pink, light blue, and orange, respectively. **B.** Promoter DNA-sequence and predicted transcription factor binding sites. Two Pax6-binding sites are displayed in purple. Candidate binding sites for ETV5 (bold), c-Jun (grey), c-Maf (green) and AP-2α (browngreen) are also shown. DNA sequence alignment displayed are: hamster (h), rat (r), and mouse (m). The TATA-box sequences are shown in yellow box.

### Statistical analysis (non-proteomics experiments)

Unpaired t test or One-way ANOVA with multiple comparisons posttest was performed using GraphPad Prism 10 for Windows (www.graphpad.com; GraphPad software) and statistical significance was considered according to p<0.001***; p<0.01**; p<0.5*. All analysis included at least 3 biological replicates.

## RESULTS

### The mouse *Crybb3* gene loss-of-function model: Deletion of the *Crybb3* promoter region

To both generate the first loss-of-function model of the mouse *Crybb3* gene and to gain basic functional information regarding its transcriptional regulation, we decided to delete its 510 bp long promoter region (-260/+250). To define the length of the “optimal” deletion, we used our earlier data showing *in vivo* “open” chromatin domain detected by ATAC-seq in E14.5 microdissected lenses (Zhao et al., 2019a). As shown in Fig. 1A, this region includes a highly conserved non-coding evolutionarily conserved region and *in vivo* Pax6-binding peak (Sun et al., 2015a). Subsequent inspection of the conserved region shows two binding sites for Pax6 flanking the canonical TATA-box, three binding sites for Etv5, one binding site for c-Maf and c-Jun, and two AP-2α binding sites (Fig.1B). Importantly, these transcription factors, i.e., Pax6, c-Maf, Etv5, and c-Jun, also bind *in vivo* and regulate the mouse αA-crystallin locus (Xie and Cvekl, 2011; Xie et al., 2016).

The *Crybb3* promoter-deleted mouse line was generated by CRISPR/Cas9-mediated genome editing (see Methods for details) and the genomic deletion was confirmed by PCR (Fig. 2A). As expected, the WT DNA generated the entire ∼800 bp promoter sequence, whereas the knockout shows the markedly reduced size of the amplicon (Fig. 2A). No survival and/or breeding issues were noted for over three years of the mice colony maintenance and experimentation. We next checked the relative expression of Crybb3 compared to GAPDH mRNAs by quantitative RT-PCR using cDNAs generated from wild type (WT), heterozygous, and homozygous cDNAs showing that the *Crybb3* knockout mice expresses very limited amounts of mutated βB3-crystallin mRNAs compared to the WT and heterozygous samples (Fig. 2B). Next, we confirmed the grossly reduced βB3-crystallin protein expression in newborn lens by immunofluorescence (Fig. 2C). The *Crybb3* knockout showed only background levels of anti-βB3-crystallin staining as well as a variable reduction in lens size reduced. Finally, major reduction/complete depletion of the βB3-crystallin proteins was determined by Western blot of whole newborn lens extract, using brain extract as a negative control, and β-actin used as a loading control (Fig. 2D).

**Figure 2.**
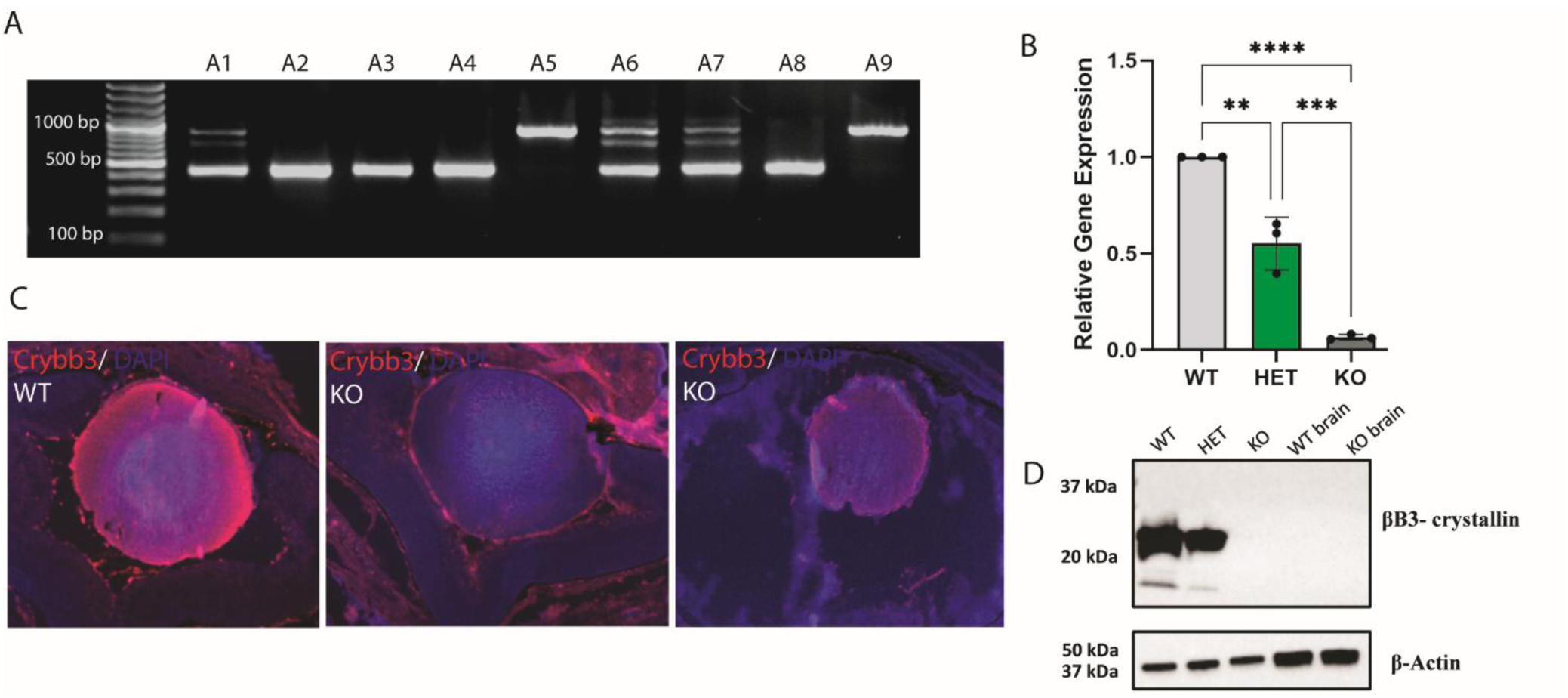
*In vivo* Deletion of the Crybb3 promoter region. **A.** PCR gel showing WT 850 bp bands and promoter excision ∼400 bp bands. A2, A3, A4, A8 display the promoter excision, A5, A9 are WT whereas A1, A6, A7 are heterozygous. **B.** Crybb3 relative gene expression by qPCR shows significant reduction in mRNA levels in the KO group. Each dot represents one biological replicate. **C.** Newborn lenses from WT, HET and KO mouse lines immunolabelled using antibodies specific for Crybb3 (red) and DAPI-counterstained. **D.** Western blot showing decreased Crybb3 protein levels in lens of Crybb3 promoter excision mice. The βB3-crystallin levels in the brain of WT and KO mouse were included as a negative control. Western blot of β-actin protein levels as a positive control in all groups analyzed. Scale bar = 100 µm.

### Lens developmental and postnatal abnormalities in *Crybb3* hetero- and homozygous lenses

Overall, the *Crybb3* homozygous knockout mice display variable lens phenotypes ranging from moderate to major gross morphological abnormalities in which the most severe phenotype is the unilateral lack of the eyecup found in approximately in 3.3% of newborn pups (Supplementary Fig.S1). Some knockout lenses already display considerably reduced sizes compared to the WT controls at birth, whereas the heterozygous lenses consistently did not show any major morphological defects. The moderate abnormalities are represented by reduced lens size only (Fig. 3). Importantly, most of the *Crybb3* homozygous knockout mice displayed moderate lens morphological abnormalities with decreased lens sizes and a smaller organelle-free zone when compared to the control group (Supplementary Fig. S1). No gross histological abnormalities were found in *Crybb3* heterozygous eyes (Supplementary Fig. S2). Finally, no retinal abnormalities were seen in either knockout or heterozygous mice (Fig. 3).

**Figure 3.**
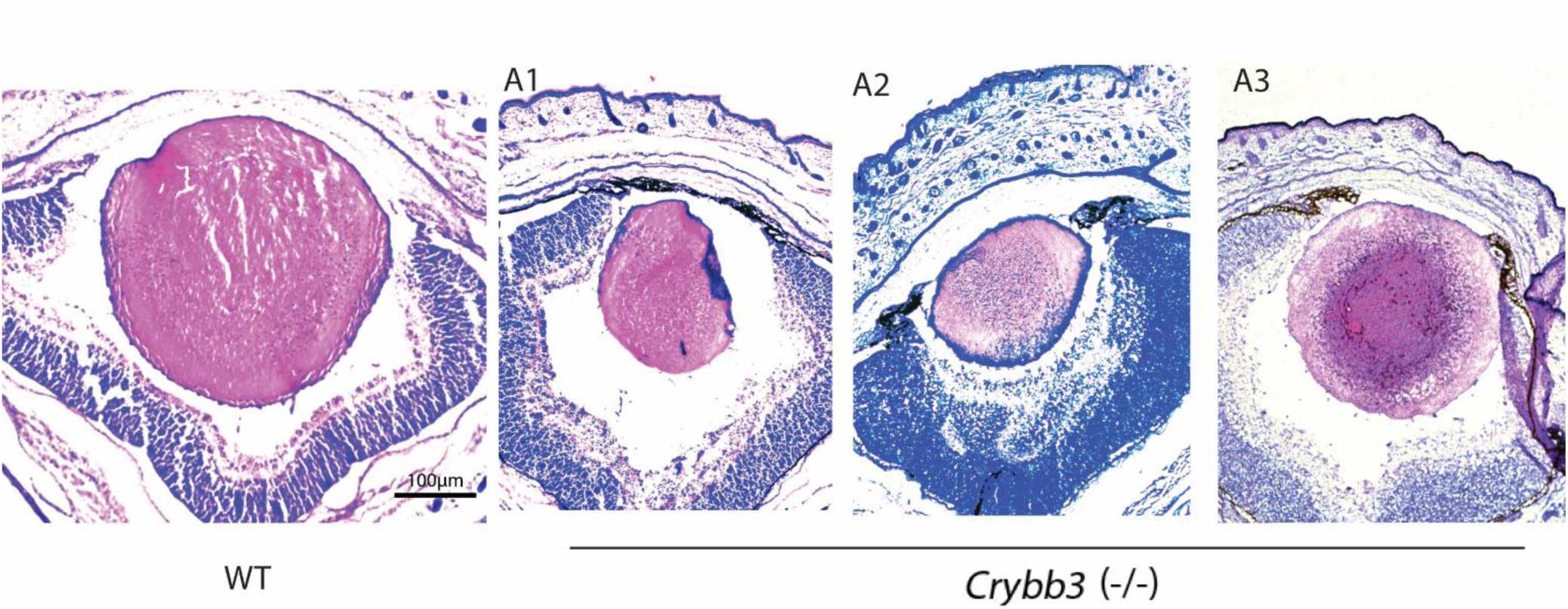
*In vivo* Deletion of the Crybb3 promoter region moderately disrupts lens morphology of newborn mice. H&E staining showing lenses morphology of wild type (WT) and three *Crybb3* knockout (A1, A2 and A3). The phenotype variability in size is observed, however, overall lens size is often reduced when compared to newborn WT.

### Proteomic studies of control and βB3-crystallin homozygous lenses

To better understand the overall impact of βB3-crystallin depletion within the protein landscape of knockout lenses, we characterized and compared four proteomics data sets from newborn, 3- and 6-week-old, and 3-month-old wild type and knockout *Crybb3* mice using isobaric labelling quantitative proteomics. The high relative abundance of crystallins in lens poses challenges to deep proteome profiling and 2,989 proteins could be quantified. We observed that small numbers of proteins were up- and down-regulated proteins (an edgeR multiple testing corrected p-value of 0.10 or smaller) in the *Crybb3* knockout mice tend to increase with age, while most of the 2,000 proteins (around 2,900) remained unchanged (Fig. 4A-D).

**Figure 4.**
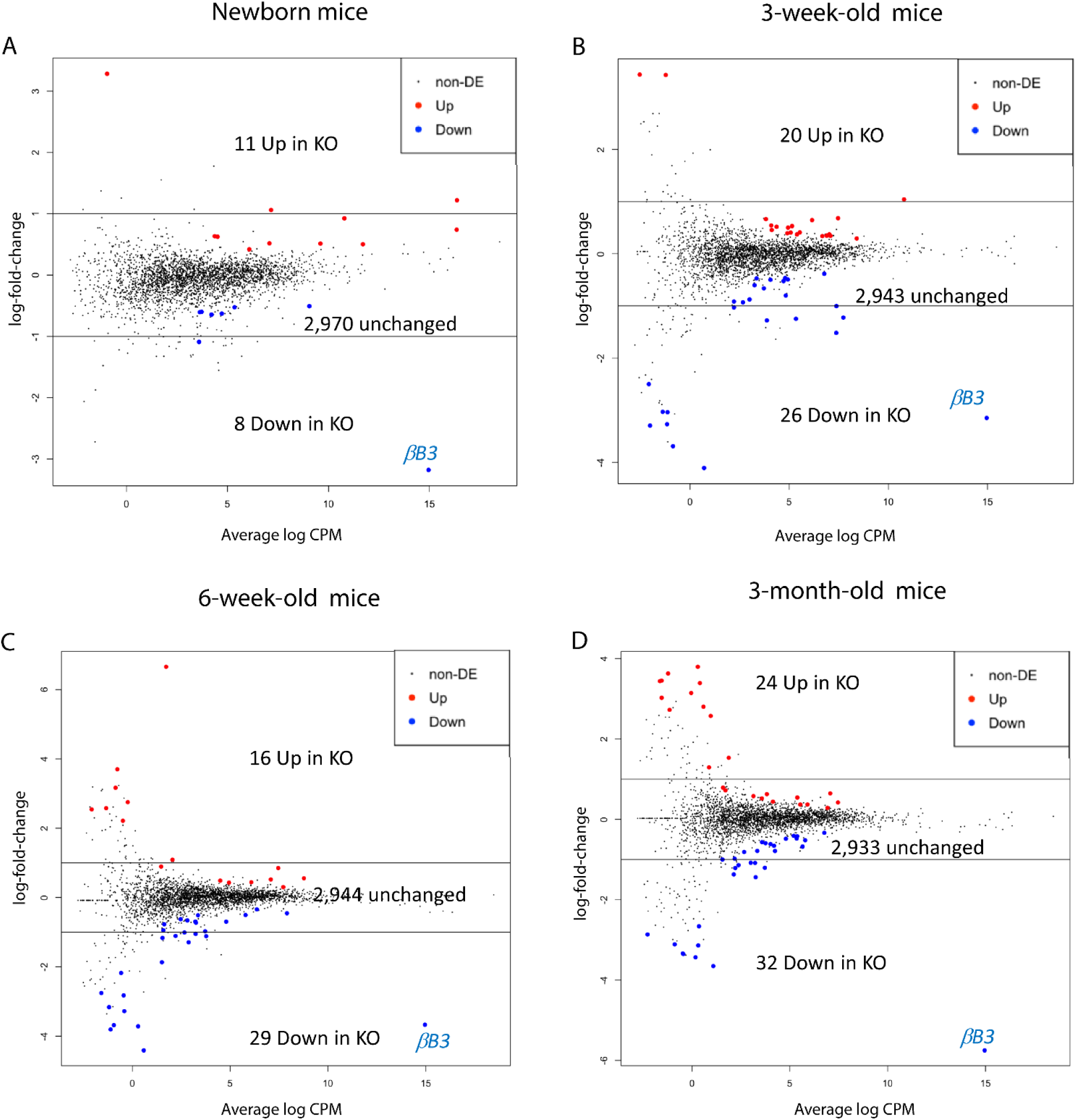
Proteomics analysis of Crybb3 promoter excision throughout lens development. Up (red) and down regulated (blue) proteins are highlighted on MA plots. The y-axis is the log ration (M) of intensities and the x-axis is the average (A) of the two group mean intensities. Unchanged proteins are in black. **A.** Newborn lenses proteins from KO and WT. 11 proteins show upregulation, whereas 8 proteins show downregulation of a total of 2,970 unchanged proteins. **B.** 3-week-old mice display 2,943 unchanged proteins compared to the control group, 20 upregulated and 26 downregulated. **C.** 6-week-old mice display 2,944 unchanged protein levels compared to the control groups whereas 16 proteins show upregulation and 29 are downregulated. **D.** 3-month-old mice display 2,933 unchanged proteins, 24 upregulated and 32 downregulated when compared to the control group. The significance cutoff for differential abundance was an edgeR Benjamini-Hochberg corrected p-value of 0.10 or less.

The increase in number of significant proteins is partly due to increasing crystallin abundances as the lens matures “pushing” the intensities of non-crystallin proteins towards the limits of quantitation, as seen in Table 1. Many of the up and down regulated proteins are on the left side (low abundance) of the MA plots, particularly for the three older ages. The union of the up- and down-regulated proteins was 131; however, 106 of these were statistically significant in only one age. This is a consequence of the dynamic range of the lens proteome and the limited linear quantitative range of TMT-based methodology. The lowest abundance proteins have fewer identified peptides and less reliable measurements resulting in inconsistent statistical significance across lens ages. There were 25 proteins statistically significant in 2 or more ages, 8 significant in 3 or more ages, and only 2 proteins statistically significant in all 4 ages.

**Table 1.** Percentages of combined crystallin TMT intensity out of total intensity for wild type and βB3 knock out mice lenses for each pooled replicate at each age. Crystallin total abundance increases by 19% between newborn and 3-week ages, by 3% between 3-week and 6-week ages, and by 1% between 6-week and 3-month ages.

| | Wild Type | | βB3 Knock Out | |
| --- | --- | --- | --- | --- |
| Age | Replicate 1 | Replicate 2 | Replicate 1 | Replicate 2 |
| Newborn | 57.5% | 67.1% | 65.1% | 68.0% |
| 3 weeks | 83.4% | 83.9% | 83.2% | 83.1% |
| 6 weeks | 87.4% | 86.8% | 86.5% | 85.1% |
| 3 months | 87.7% | 88.6% | 88.0% | 87.1% |

The small number of up- or down-regulated proteins between wild type and knock out and their modest fold changes suggest that the mouse lens can adapt to loss of βB3-crystallin; nevertheless, its absence is detrimental for normal lens function. Data from figures 4B-D coupled the consistent relative crystallin abundances in Table 1 for the three most advanced ages suggest a stable readjustment of crystallin subunits in the knock-out lenses. Some differences in crystallin abundances were present in newborn lenses (Fig. 4A and Table 1) but not present in the older lenses suggesting this readjustment may occur early in development.

One of the two proteins significantly downregulated in all ages was βB3-crystallin, confirming the successful promoter excision model (βB3-crystallin is highlighted in blue in Figs. 4A-D). TMT pro tags, with naturally occurring C13 isotopes and N14 and C12 impurities in the heavy N15 and C13 purified isotopes, result in intensities for each sample having roughly 8% of their signal coming from adjacent m/z tags. This limits the dynamic range of TMT intensity differences and lower limits of tag intensities measurements (on the order of 5-10% of adjacent m/z intensities). The relative intensities for βB3-crystallin in the knock-out samples were 7-8% of the intensities of the wild type samples, consistent with little or no βB3-crystallin present in the knock-out lenses.

More detailed inspection of dysregulated proteins found that both βB3-crystallin and a regulatory subunit of the BAF complexes Smarcc1/Baf155 were downregulated at all four time points analyzed (Table 2). Six additional proteins: 10 kDa heat shock protein (Hspe1), PDZ and LIM domain protein 1 (Pdlim1), Aspartate aminotransferase (Ast/Got), U6 snRNA-associated Sm-like protein LSm7 (Lsm7), RNA helicase (Ddx23), and Acyl-CoA dehydrogenase family member (Acad11) were downregulated in at least three of four-time comparisons (Table 2). Taken together, the present data show increasing perturbation of the lens proteome from earlier to advanced time points.

**Table 2.**
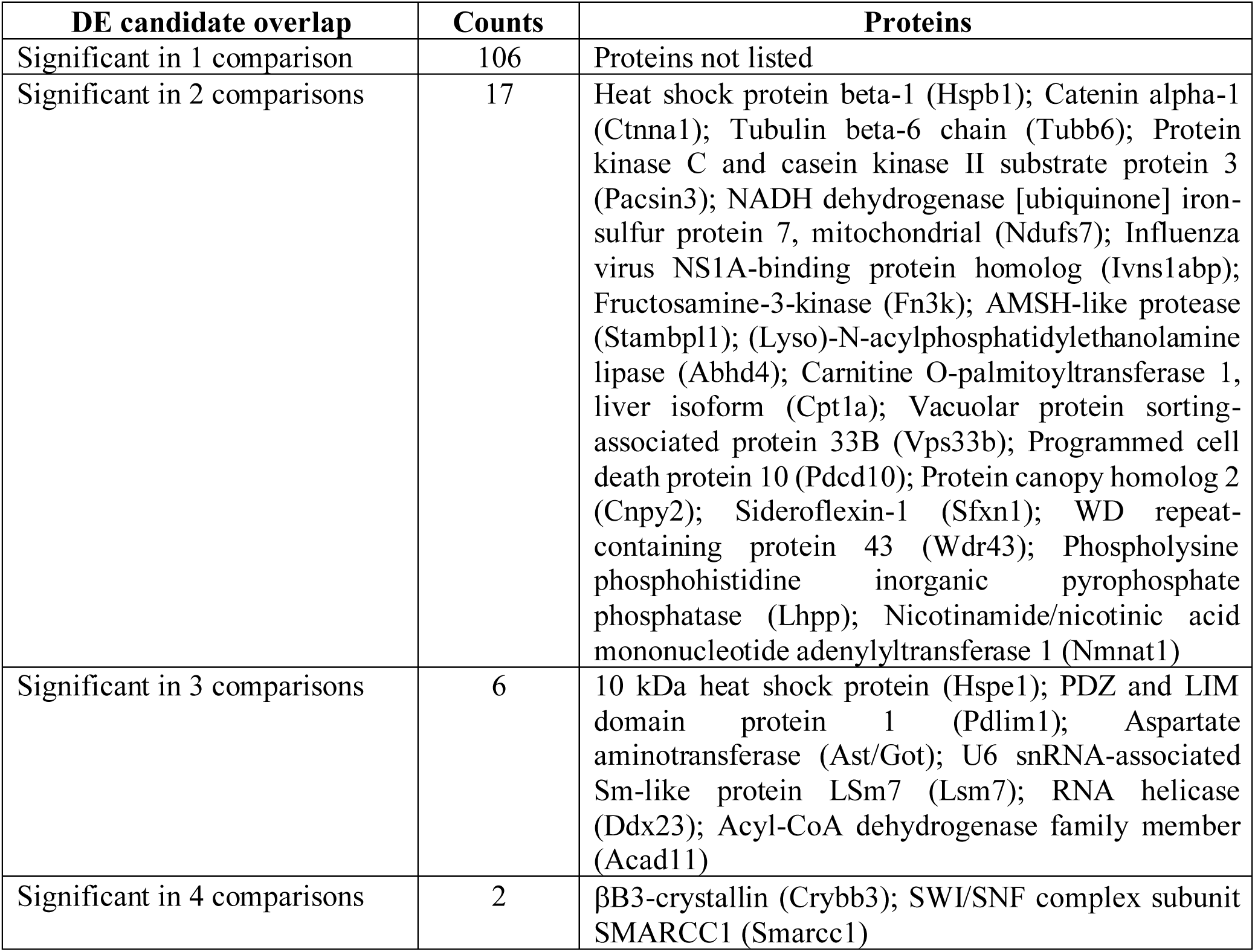
Overlap of proteins significantly changed in relative abundance in 1, 2, 3, or 4 age group comparisons. A multiple testing adjusted edgeR significance cutoff of 0.10 was used. Less reliable and low abundance proteins are more likely to be significant in only one comparison. Ranking protein abundance by average intensity of all 16 samples and excluding the bottom 25% of the proteins reduced the numbers in second column to 67, 12, 6, and 2, respectively. Candidates seen in multiple comparisons may have more biological significance. Proteins in the third column are listed by decreasing relative abundance. See Supplemental Table 1 for other protein candidates.

### Activation of the Crybb3 promoter by FGF2 in cultured lens cells

FGF signaling plays a major role in regulation of lens development and crystallin gene expression (see McAvoy et al., 2017; Makrides et al., 2022). We therefore asked if the mouse βB3-crystallin promoter (-250/+210) was responsive to FGF2 in cultured primary chicken lens epithelial cells. As shown in Fig. 5, the WT promoter drove expression of the EGF reporter in a manner enhanced by FGF2 treatment. Interestingly, two mutations of the βB3-crystallin promoter that lack the Pax6 binding site had higher activities compared to the WT construct in either the absence or presence of FGF, in agreement with Pax6 functioning as transcriptional repressor of the βB3-crystallin promoter (Fig. 5A-C) and shown earlier for the βB1-crystallin reporter construct (Duncan et al., 1998).

**Figure 5.**
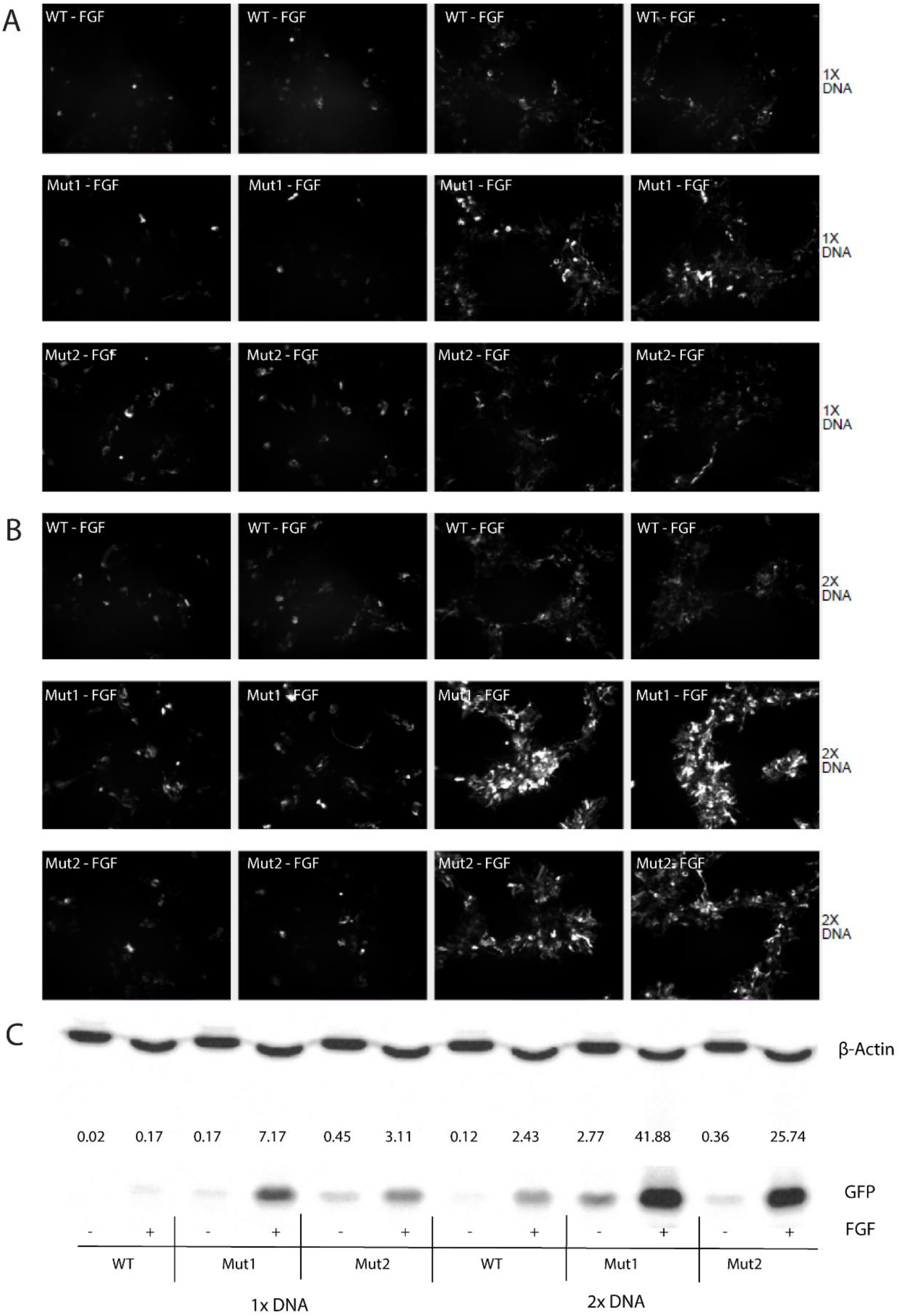
Up-regulation of the βB3-crystallin promoter by FGF. EGFP+ primary embryonic lens epithelial cells from E10 chick embryo lenses were culture with FGF-2. Mutant cells lacking one or two Pax6 binding sites (M1 and M2 respectively) were transfected with 1X and 2X reporter plasmid DNA (A-B). C. M1 and M2 cell lines display significantly higher proliferation in response to FGF-2 when compared to WT cell line. Β-actin control bands (upper); EGFP bands in WT, M1 and M2 cell lines (bottom).

## DISCUSSION

The present study generates the first loss-of-function model of the βB3-crystallin gene which encodes highly lens-preferred protein. Heterozygous mutations in human *CRYBB3* gene cause early congenital cataracts (Riazuddin et al., 2005; Jiao et al., 2016; Reis et al., 2013; Li et al., 2016) while aging processes within the lens impact structural integrity and native conformations of individual crystallins leading to age-related cataracts (see Benedek, 1997; Bloemendal, et al., 2004; Lampi et al., 2014). The early lens abnormalities following depletion of the mouse βB3-crystallin contrast with depletion of the comparable βB2-crystallin that resulted in normal lenses in very young animals with cortical cataracts only developing after several months (Zhang et al., 2008). This supported that unlike βB2-crystallin, βB3-crystallin may play its most important role in early lens development. The decrease in βB3-crystallin during aging in lens is also consistent with this hypothesis (Ueda et al., 2002, Halverson et al., 2024).

CRISPR-Cas9 genome editing technology facilitates the generation of rodent models for the analysis of human missense and nonsense mutations, either naturally occurring or selected for study based on 3D-models and sequence analysis. Given the lack of earlier rodent models, we decided to generate somatic depletion of the βB3-crystallin by deleting its 510 bp promoter region as this approach can be followed by subtle deletion/mutations of individual *cis*-regulatory sites (see below for details). Our Western blot analyses showed absence of the protein (Fig. 2D); however, the qRT-PCR data identified residual signals (Fig. 2B) that could be caused by alternate minor transcripts without any coding capacity. The proteomic data reflected the reduced βB3-crystallin protein abundances in the knockout lenses, which were at the lower limits of measurement.

The initial phenotypic analyses were focused on lens/eye morphological changes over the course of three months. As expected, variability of the phenotype was observed in newborn lens ranging from light (reduced lens size measurements), moderate (reduced lens size measurements and reduced organelle free zone area) and severe (unilateral eye cup absence) phenotypes (Supplementary Fig. S1). However, most animals analyzed show moderate lens morphological abnormalities, marked by smaller overall lens size coupled with smaller organelle free zone at the lens center, when compared to WT eyes.

The quantitative proteomics data from newborn, 3- and 6-week-old, and 3-month-old wild type and knockout *Crybb3* mice showed surprisingly small effects from loss of a major mouse lens crystallin. There were few statistically significant protein abundance differences at any of the 4 ages. The few up or down regulated proteins (except for very low abundance proteins) mostly had changes of less than 2-fold. The relative fractions of total crystallin intensity out of the total proteome intensity was also essentially unchanged between wild type and knock out at all ages (Table 1). Interestingly, αA- and βB2-crystallins are up regulated in newborn knock-out lenses perhaps playing a role in the readjustment of crystallins. The newborn lens may have differences in response to loss of βB3-crystallin; however, more replicates and additional time points around this active stage of development are needed.

It is important to state here that depletion of the βB3-crystallin can cause different phenotypes compared to the missense and nonsense heterozygous or homozygous mutations, and their impacts can be different between human and mouse. Given major progress in generation of human lenses from induced pluripotent stem (iPS) cells (Li et al., 2024), it is now possible to generate a series of *CRYBB3* cell lines carrying both hetero- and homozygous cataract mutations within an isogenic genetic background (see Cvekl and Camerino, 2022).

Smarcc1/Baf155 is a regulatory subunit of the highly polymorphic BAF complexes (see Hodges et al., 2016; Centore et al., 2020) containing two Baf155, Baf155/Baf170, or two Baf170 subunits. Transfer of Brg1, a catalytical subunit of the BAF complexes, into the lens cytoplasm in terminally differentiating lens fiber cells (Limi et al., 2018) and role in protein translation outside of the lens have been recently shown (Ulicna et al., 2022; Nguyen et al., 2024). Depletion of Brg1, a catalytical subunit of the BAF complexes, causes lens developmental abnormalities with variable phenotypes seen at E13.5 embryos (He et al., 2010). Thus, it is possible that reduction of Smarcc1/Baf155 due to loss of βB3-crystallin is responsible for variability of early lens abnormalities found here.

The results of deletion of the Crybb3 promoter fragment that we report here, coupled with evidence that the Crybb3 promoter contains individual binding sites for Pax6, c-Maf, Etv5, c-Jun, and AP-2a, can be followed by deletions/mutations of these sites to better understand the *cis*-regulatory grammar of the *Crybb*3 gene and compare it to the regulation of the βB2-crystallin gene outside of the lens (Gao et al., 2014; Heermann et al., 2019; Li et al., 2021; Bauer et al., 2024). Our findings that point mutagenesis of either Pax6-binding site in the Crybb3 promoter increases the basal activity of the reporter compared to the control (Fig. 5) is consistent with Pax6 acting as a repressor of the highly related βB1-crystallin (Duncan et al., 1998). In addition, our recent single cell RNA-seq studies have shown that Crybb3 mRNAs are already expressed in the invaginating lens placodes/pits of the E10.5 mouse embryos. This is prior to the formation of primary lens fiber cells and may indicate a novel transcriptional mechanism for the onset of Crybb3 transcription (Camerino et al., 2025 in preparation) directly linked to accumulation of spliced Crybb3 mRNA in the early lens fiber cell nuclei prior their transfer into the cytoplasm and translation in lens fiber cells (Limi et al., 2019).

In conclusion, the present data provide in-depth analysis of control and βB3-crystallin depleted lens proteomes using the C57Bl6 mouse strain and time-course pattern of significantly changed proteins will serve for comparative purposes between other mouse models and iPS-derived human 3D-lentoid cultures with particular focus on the analysis of cataract-causing CRYBB3 missense mutations.

## Supporting information

Rayee et al 2024 Supplemental File

## Abbreviations

H&E: Hematoxylin and eosin
iPS: induced pluripotent stem
OCT: optimal cutting temperature
WT: wild type

## ACKNOWLEDGEMENTS

We thank Dr. Yongwei Zhang and Dr. Winnfried Edelmann (Einstein Gene Modification core facility) for the generation of the mouse model. Funding: R01 EY014237 (AC). Mass spectrometric analysis was performed by the OHSU Proteomics Shared Resource with partial support from NIH core grants P30EY010572, P30CA069533, and S10OD012246.

## FIGURE LEGENDS

**Supplementary Figure S1: Phenotype variability in KO mice lacking Crybb3 promoter sequence. A.** P0 and 3-month-old mice displaying complete left eye absence. **B.** KO mice display a variable lens morphology spectrum from light to severe (eye absence) phenotype. Most of the animals analyzed display moderate lens morphological abnormalities. N= 15 KO analyzed animals from independent litters at P0 stage compared to WT. Parameters measured: lens area relative to control; organelle-free zone absence.

**Supplementary Figure S2: WT, heterozygous and knockout newborn lenses morphology. Supplementary Table S1. Proteomics studies of lens after Crybb3 promoter excision**.

