## Supplementary figures and images for "Analysis of mouse lens morphological and proteomic abnormalities following depletion of βB3-crystallin"

### Rayee et al 2024 Supplemental File

Supplementary Fig. 1

a.

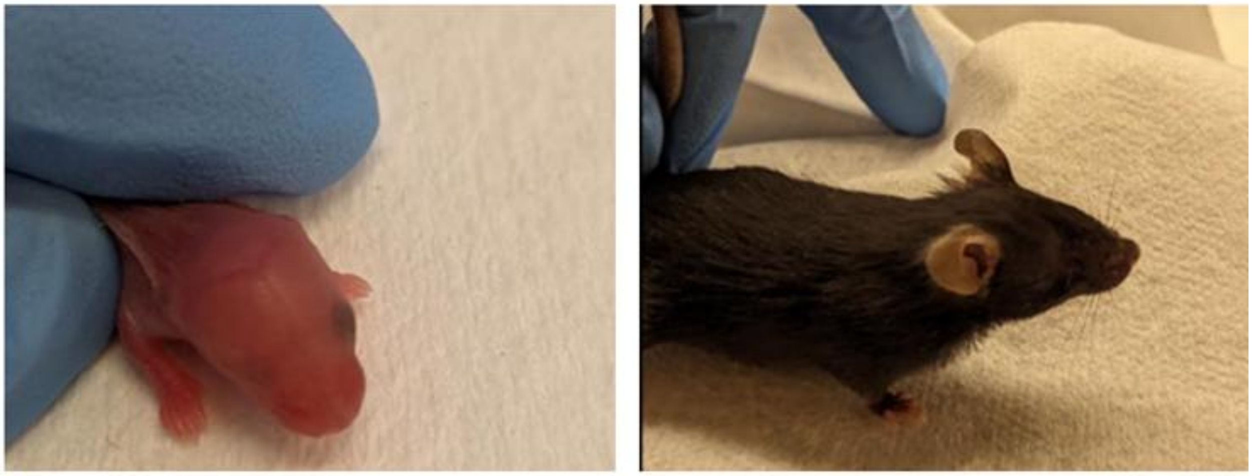

b.

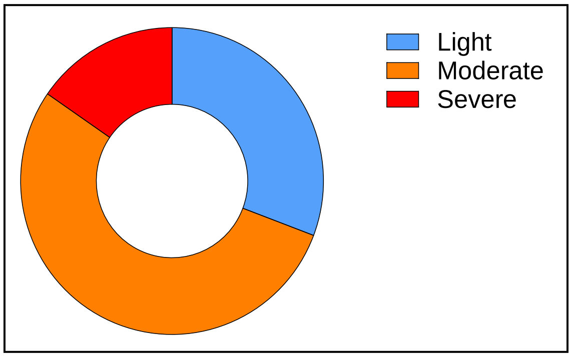


Supplementary Fig. 2

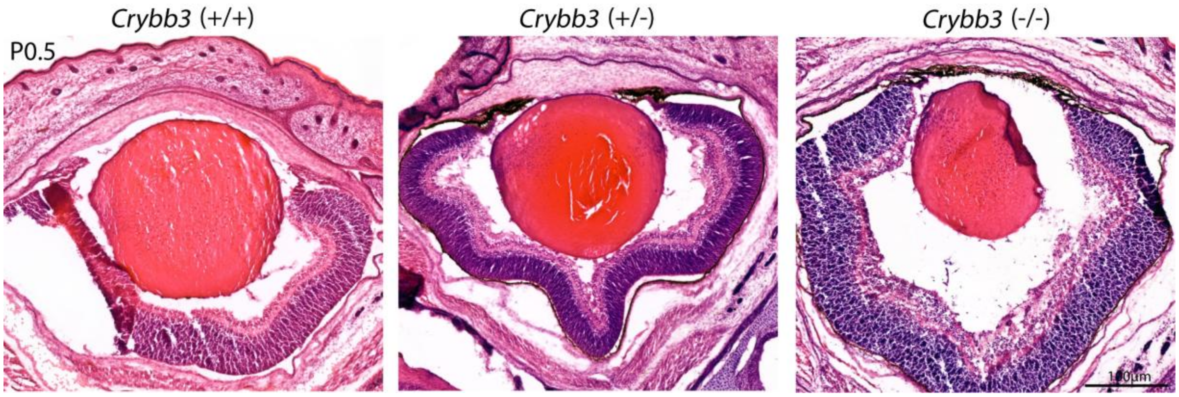
